# FOXP1 differentially regulates the development of murine vasopressin and oxytocin magnocellular neurons

**DOI:** 10.1101/2025.06.10.658830

**Authors:** Jari Berkhout, Sophie Trender, Quirin Krabichler, Yuval Podpecan, Felix Franke, Tim Schubert, Peter Burbach, Valery Grinevich, Roger Adan, Henning Fröhlich, Ferdinand Althammer, Onno Meijer, Ahmed Mahfouz

## Abstract

The neuropeptides arginine vasopressin (AVP) and oxytocin (OXT) are closely related. As neurohormones, AVP and OXT are mainly produced in magnocellular neurons (MCNs) located in the hypothalamus. Development of both neuron types requires coordinated expression of transcription factors OTP, SIM1, ARNT2 and POU3F2. However, the exact transcription factors involved in the differential differentiation of the AVP and OXT lineages are yet unknown. We used a publicly available single-cell RNA-sequencing dataset of the developing mouse hypothalamus to identify gene regulatory networks linked to AVP and OXT neuronal differentiation. We identified RORA, EBF3, FOXP1, FOXP2, and BCL11B as transcription factors with possible relevance for *Avp* and *Oxt* MCN divergence. We then modeled developmental gene expression dynamics using computational cell fate mapping. This revealed enrichment of EBF3 and BCL11B in the *Avp* lineage, while FOXP1 and FOXP2 are enriched in the *Oxt* lineage. Next, *in silico* analysis of *Avp* and *Oxt* promoters found predicted binding sites for FOXP1 and FOXP2 in the *Oxt* promoter, suggesting a role in *Oxt* MCN differentiation. Finally, we validated the role of one candidate (FOXP1) with a heterozygous knockout mouse line. Compared to wild-type littermates, we find decreased AVP and OXT neuron abundance, with OXT neurons disproportionally affected.

## Introduction

A long-standing open question in neuroendocrinology is the origin of the transcriptional differences between the hypothalamic magnocellular neuron (MCN) types. These neurons are located in the hypothalamic paraventricular nucleus (PVN) and supraoptic nucleus (SON), where they produce the neuropeptides arginine vasopressin (AVP) and oxytocin (OXT). Structurally, AVP and OXT are closely related nonapeptides, differing at only two amino acids. The genes encoding for AVP and OXT are only 10 kb apart on the human genome, and thought to be the result of a gene duplication event around the origin of vertebrates ^1,2^. Despite their evolutionary relatedness, AVP and OXT differ in hormonal function: AVP regulates arterial blood pressure and water homeostasis^3^, while the best known function of OXT is the induction of parturition^4^ and lactation^5^. As systemic hormones, AVP and OXT are released from MCNs projections to the posterior pituitary. Despite their similarities, AVP and OXT are predominantly produced in neuropeptide- specific MCNs^6^.

AVP and OXT MCNs are clearly distinct, yet similar in some respects. A common feature is the known developmental trajectories of both lineages. Development of both AVP and OXT MCNs necessitates expression of transcription factors (TFs) OTP, SIM1, ARNT2 and POU3F2 ^7–12^. Also, it has been known for some time that AVP and OXT MCNs express mRNA for their counterpart peptides, though at levels differing orders of magnitude^13,14^. Recently, we corroborated these findings with a single-cell transcriptomic atlas of the PVN, and found the AVP and OXT MCNs to be highly transcriptionally correlated, yet also distinct^15^.

The evolutionary relatedness, high level of transcriptional similarity and largely overlapping ontogeny implies a common developmental progenitor. Yet, the presumed transcription factor(s) involved in the divergence of the AVP and OXT lineages have not been described yet. While several candidate transcription factors have been proposed to potentially drive differential expression of *AVP* and *OXT*^16^, none have been conclusively established to do so. In this work, we aim to answer this question using publicly available single-cell transcriptomics data, and to validate one of the identified candidate TFs *in vivo*.

## Results

### Publicly available single-cell RNAseq reveals all developmental stages of MCNs

We used the developmental single-cell dataset from Romanov *et al*. (2020)^17^, comprising of mouse hypothalamus tissue at several developmental timepoints: embryonic days 15 and 17 (E15, E17), and post-natal days 0, 2, 10 and 23 (P0, P2, P10, P23; Fig. 1A). A subset of the dataset was taken based on annotations as defined by Romanov *et al*. (2020)^17^, to only include MCNs. The subset was then integrated to eliminate internal batch effects, and subsequently clustered based on the integrated embedding.

**Figure 1.**
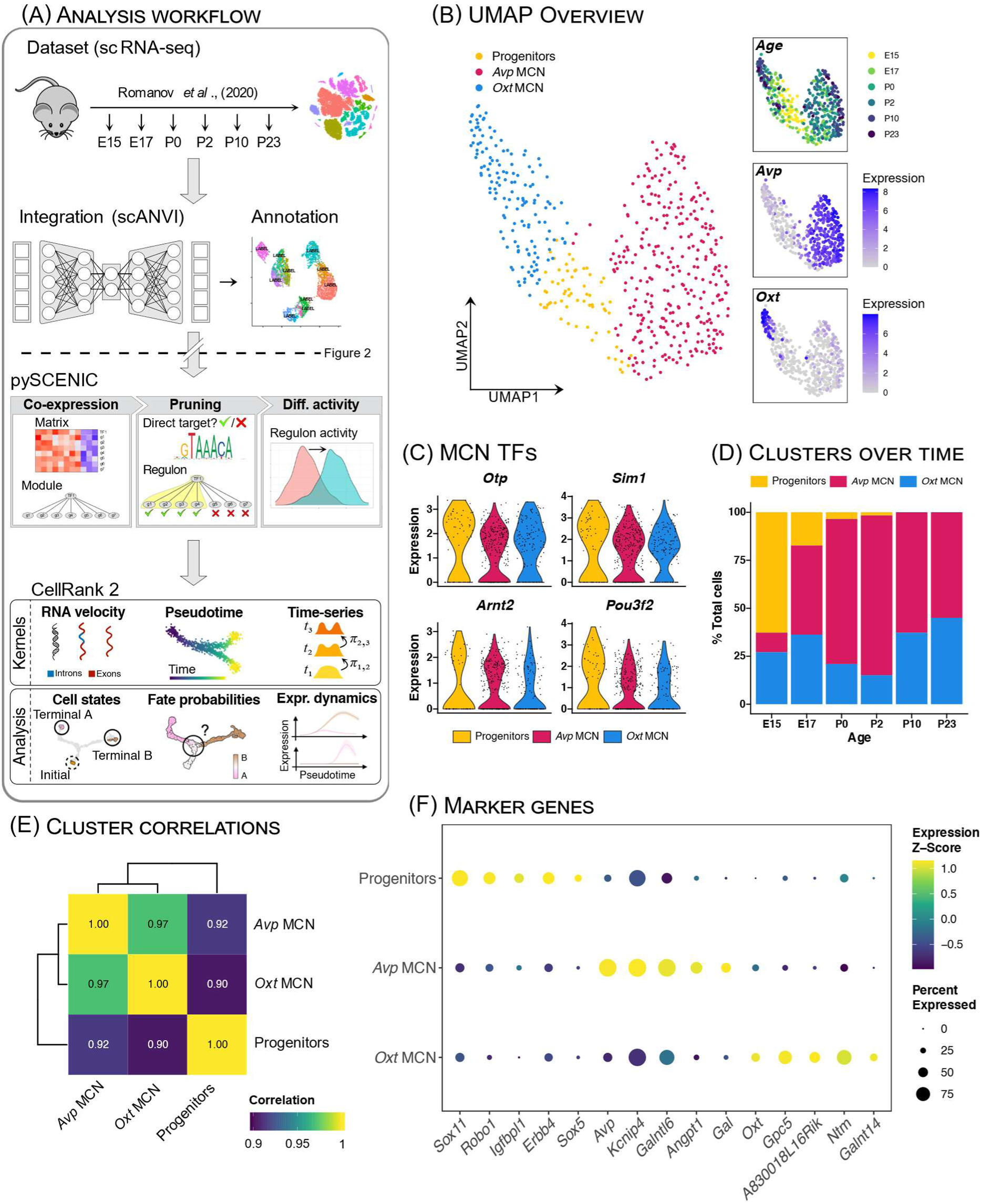
Developmental single-cell RNA-seq (scRNA-seq) of magnocellular neurons (MCN) reveals three distinct clusters **A.** Workflow of the analysis presented in this Fig. 1 and Fig. 2. The developmental mouse dataset from Romanov *et al.* (2020)^17^ was integrated with scANVI^19^, and then clustered and annotated. Subsequently, the pySCENIC^18^ pipeline was performed to infer gene regulatory networks. Finally, the CellRank 2 pipeline^20^ was applied to infer cell fate probabilities. **B.** (Left) Uniform Manifold Approximation and Projection (UMAP) showing the clusters as annotated after integration. (Right) UMAPs with coloring representing sampling timepoints (Age), *Avp* expression and *Oxt* expression. **C.** Violin plots of the expression levels for known MCN-expressed transcription factors (TFs) *Otp, Sim1, Arnt2,* and *Pou3f2*. **D.** Stacked bar plots of the proportion of cells annotated as each cluster over the different sampling timepoints. **E.** Heatmap of the correlation coefficients between pseudobulk expression profiles for each cluster in the dataset. All clusters are highly correlated, coloring is scaled to show mutual differences in correlations. F. Dotplot of the normalized expression levels for cluster-enriched gene markers. Coloring represents Z-scaled expression levels, dot size represents percentage expression of a marker within the respective cluster.

We identified three clusters of MCNs: progenitors, *Avp* MCNs and *Oxt* MCNs (Fig. 1B). The *Avp* and *Oxt* MCNs strongly expressed *Avp* and *Oxt*, respectively, while the progenitors did not express either neuropeptide (Fig. 1B). All three clusters expressed high levels of the currently known MCN TFs *Otp, Sim1, Arnt2,* and *Pou3f2* (Fig. 1C). As expected, the progenitor cluster was most abundant at the earlier timepoints, diminishing in abundance over time (Fig. 1D). All clusters were highly correlated to each other, with the progenitor cluster being relatively most distinct from both *Avp* MCNs and *Oxt* MCNs (Fig. 1E). Progenitors were characterized by their high relative expression of *Sox11, Robo1, Igfbpl1, Erbb4,* and *Sox5*. *Avp* MCNs were characterized by their expression of *Avp, Kcnip4, Galntl6, Angpt1,* and *Gal.* Finally, *Oxt* MCNs were characterized by expression of *Oxt, Gpc5, A830018L16Rik, Ntm,* and *Galnt14* (Fig. 1F).

### RORA, EBF3, FOXP1, FOXP2, and BCL11B are candidate TFs for diverging differentiation

Next, we used pySCENIC^18^ to identify candidate gene regulatory processes underlying Avp-Oxt differentiation. pySCENIC infers gene regulatory networks (GRNs) centered around TFs – also known as regulons –through co-expression analysis paired with transcription factor binding site (TFBS) enrichment analysis (Fig. 1A). A regulon can be based on either positively correlating components or negatively correlating components, resulting in activating and repressive regulons, respectively. An activity score is then assigned for each regulon in each cell, which we used to calculate differentially active regulons between the *Avp* and *Oxt* clusters (Fig. 2A). To narrow down our selection, regulons were filtered out if the associated TF was not a significantly differentially expressed gene between the *Avp* and *Oxt* clusters.

**Figure 2.**
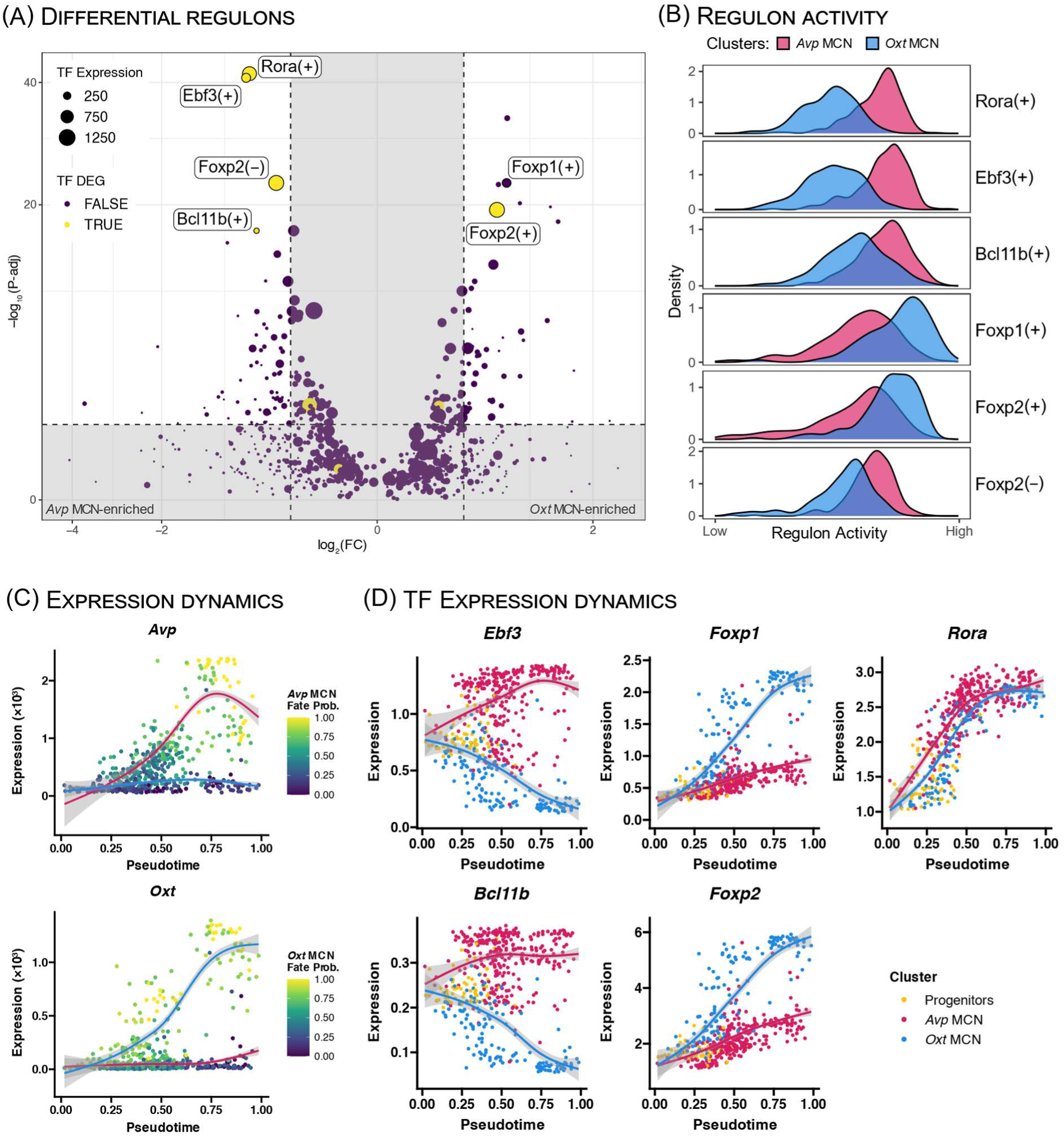
Rora, Ebf3, Foxp1, Foxp2, and Bcl11b are candidate TFs for diverging *Avp* vs *Oxt* MCN differentiation. **A.** Volcano plots with differential regulons activity scores in *Avp* clusters (left) versus *Oxt* clusters (left). Activating regulons are denoted with a plus sign, repressive regulons with a minus sign. Dot size represents the gene expression levels of the TF defining the regulon. Dot colors represent if the gene expression of the TF defining the regulon is differential between clusters. **B.** Density plots of the regulon activity score in the *Avp* and *Oxt* MCN clusters, aggegrated over all timepoints. **C.** Denoised gene expression dynamics of *Avp* and *Oxt* over pseudotime. Dot colors show cell fate probabilities for differentiation towards the *Avp* MCN fate (upper) or *Oxt* MCN fate (lower). Curves are fitted using a generalized additive model weighted by the fate probability odds ratios. Curve colors correspond to coloring used in Fig. 1B. **D.** Denoised gene expression dynamics of *Rora*, *Ebf3*, *Foxp1*, *Foxp2*, and *Bcl11b* over pseudotime. Dot colors clustering assigned after integration. Curves are fitted using a generalized additive model weighted by the fate probability odds ratios. Dot and curve colors correspond to coloring used in **Fig 1B**.

With these criteria, the activating regulons Rora(+), Ebf3(+), Foxp2(+) and Bcl11b(+) were included, as well as the repressive regulon Foxp2(–). In addition, the differentially active regulon Foxp1(+) was included despite lack of differential *Foxp1* gene expression. Although not differentially expressed between *Avp and Oxt clusters*, *Foxp1* could be biologically relevant and might in fact drive the observed differential expression of the Foxp2 regulons through its well-described ability for heterodimerization with *Foxp2*^21,22^. From the selected regulons, Rora(+), Ebf3(+), Bcl11b(+) and Foxp2(–) were found enriched in *Avp* MCNs, while Foxp1(+) and Foxp2(+) were enriched in *Oxt* MCNs (Fig. 2B).

Of note, this differential activity analysis does not account for temporal dynamics. To interrogate the temporal dynamics of the resulting regulons, we applied CellRank2, utilizing RNA velocity^23^, pseudotime^24^, and optimal-transport analysis^25^ to infer cell fate probabilities (see Methods). With these cell fate probabilities, we then visualized the expression dynamics of genes throughout pseudotime (Fig. 2A).

Analyzing the expression dynamics for *Rora, Ebf3, Foxp1, Foxp2,* and *Bcl11b* revealed the expression pattern for *Rora* to be highly similar between *Avp* and *Oxt* clusters (Fig. 2B). Both *Ebf3* and *Bcl11b* expression increase in *Avp* MCNs and decrease in *Oxt* MCNs (Fig. 2B). Finally, the expression patterns for *Foxp1* and *Foxp2* were found to be highly similar to each other, both increasing in *Avp* MCNs as well as *Oxt* MCNs, albeit at a considerably faster rate in the latter (Fig. 2B). A key difference is the overall expression levels of these factors, which were higher for *Foxp2* than for *Foxp1.* This lower *Foxp1* expression and subsequent smaller absolute difference between populations may also explain the observed lack of significant *Foxp1* expression difference between *Avp* and *Oxt* MCNs. Notably, the observed expression patterns for both *Foxp1* and *Foxp2* are very similar to the expression pattern for *Oxt*.

### Promoters for *Avp* and *Oxt* contain bindings sites for RORA, FOXP1, and FOXP2

We then investigated if binding sites for the identified TFs exist within the *Avp* or *Oxt* promoters to directly regulate these neuropeptides. Previous work on the mouse *Avp* and *Oxt* promoters defined regions in both promoters that are necessary for cell-type specific expression^16,26,27^. Within these specificity-conferring regions (SCRs), we predicted the presence of TFBSs for candidate TFs (RORA, EBF3, FOXP1, FOXP2, and BCL11B) using their corresponding motifs from JASPAR^28^. The *Avp* SCR contained a single TFBS for RORA (Fig. 3A). The *Oxt* SCR contained several TFBSs for RORA, and a single TFBS for both FOXP1, and FOXP2 (Fig. 3B). These findings align with Gainer (2012)^16^, who reported predicted TFBSs in the *Oxt* SCR for RORA and FOXO1, a TF with a motif highly similar to both FOXP1 and FOXP2.

**Figure 3.**
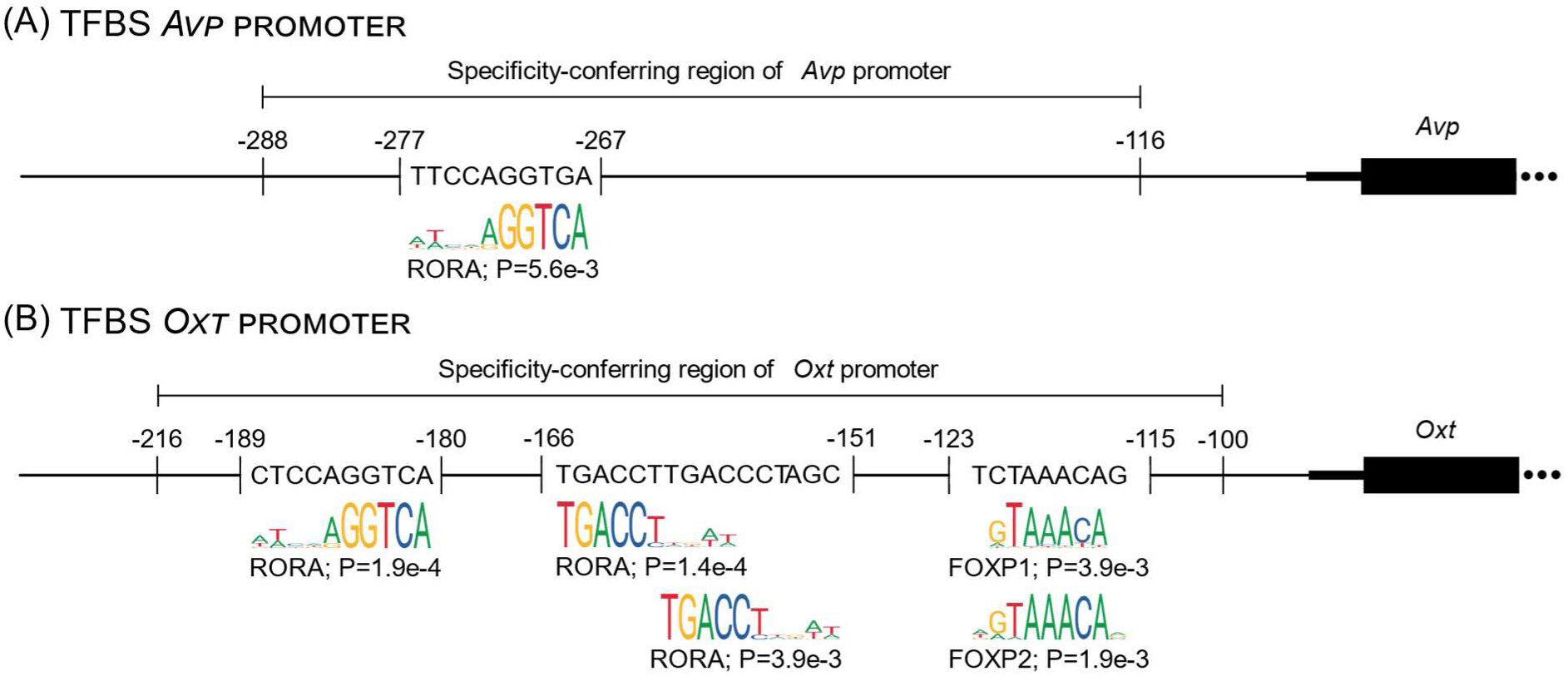
Crucial *Avp* and *Oxt* mouse promoter loci contain transcription factor binding sites (TFBSs) for RORA, FOXP1, and FOXP2 **A.** Schematic representation of the mouse *Avp* locus, with a focus on the specificity-conferring region (SCR) as defined by Ponzio *et al.* (2012)^26^. Predicted TFBSs are shown including coordinates and the aligned JASPAR motifs. Coordinate placement not to scale. Schematic representation of the mouse *Oxt* locus, with a focus on the specificity-conferring region (SCR) as defined by Fields *et al.* (2012)^27^. Predicted TFBSs are shown including coordinates and the aligned JASPAR motifs. Coordinate placement not to scale.

### *Foxp1*^+/-^ mice have disproportionally fewer OXT neurons than wild-type littermates at P10

Our results so far strongly suggest that FOXP1 and/or FOXP2 play a role in Avp-Oxt MCN differentiation by regulating *Oxt* but not *Avp* MCNs. For validation of this hypothesis in a physiologically relevant context, we opted to analyze whole hypothalami of *Foxp1* heterozygous knockout mice (*Foxp1^+/-^*), the only model available to us. Homozygous knockouts are embryonically lethal^29^, and *Foxp1*^+/-^ mice have previously been used to study the effects of *Foxp1* haploinsufficiency in the brain^30,31^. As such, PVN- and SON-containing brain sections from P10 male *Foxp1*^+/-^ mice (n=7) and wild-type (WT; n=7) littermates were stained for AVP and OXT, and subsequently imaged using confocal microscopy. The resulting 3D images were fully quantified to measure AVP and OXT neuron abundance, as previously described^32^. We hypothesized the heterozygous knockout would induce a selective reduction in OXT neurons compared to WT mice.

In the PVNs of *Foxp1*^+/-^ mice, fewer OXT neurons were observed than in WT (Fig. 4A; *t* = 5.3; P = 2.8e-4). A more modest decrease was observed for AVP neurons (Fig. 4B; *t* = 2.8; P = 0.017). Further, the ratio between OXT and AVP neurons was decreased significantly in the *Foxp1*^+/-^ mice (Fig. 4C; *t* = 3.2; P = 0.008), indicating that the partial loss of FOXP1 disproportionally affected OXT neurons.

**Figure 4.**
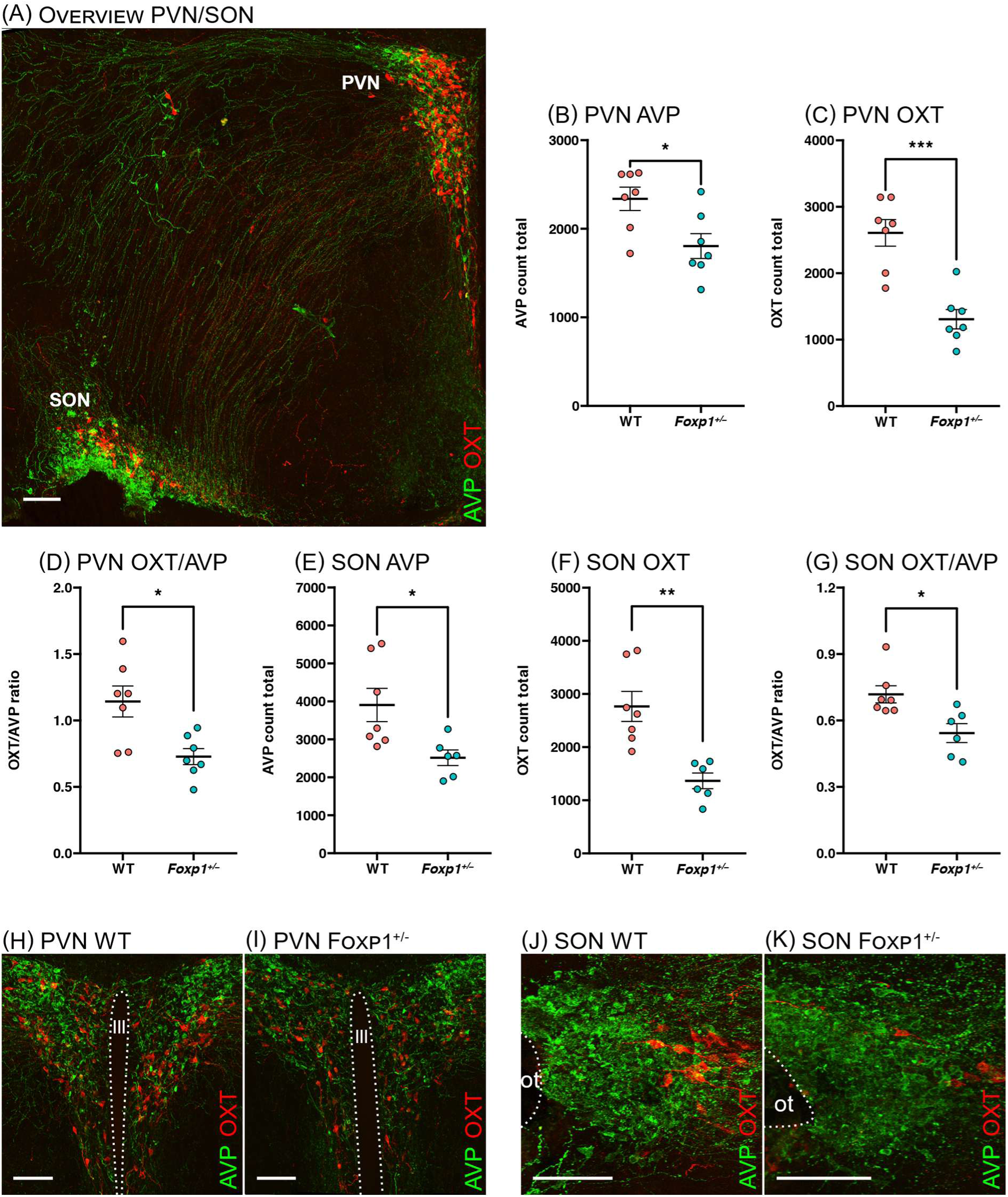
*Foxp1*^+/-^ mice have disproportionally fewer OXT neurons in both the paraventricular nucleus (PVN) and supraoptic nucleus (SON) at P10. **A.** Overview image of the mouse hypothalamus at P10, stained for AVP (green) and OXT (red). Scale bar 100 µm. **B.** Dot plot of total observed OXT neurons in the whole PVN for *Foxp1*^+/-^ mice compared to wild-type (WT) littermates. **C.** Dot plot of total observed AVP neurons in the whole PVN for *Foxp1*^+/-^ mice compared to WT littermates. **D.** Dot plot of the ratio of observed OXT/AVP neurons in the whole PVN for *Foxp1*^+/-^ mice compared to WT littermates. **E.** Dot plot of total observed OXT neurons in the whole SON for *Foxp1*^+/-^ mice compared to WT littermates. **F.** Dot plot of total observed AVP neurons in the whole SON for *Foxp1*^+/-^ mice compared to WT littermates. **G.** Dot plot of the ratio of observed OXT/AVP neurons in the whole SON for *Foxp1*^+/-^ mice compared to WT littermates. **H.** Representative image of a WT PVN, stained for AVP (green) and OXT (red). Scale bar 100 µm. **I.** Representative image of a *Foxp1*^+/-^ PVN, stained for AVP (green) and OXT (red) Scale bar 100 µm. **J.** Representative image of a WT SON, stained for AVP (green) and OXT (red). Scale bar 100 µm. **K.** Representative image of a *Foxp1*^+/-^ SON, stained for AVP (green) and OXT (red). Scale bar 100 µm. Asterisks indicate statistically significant differences as determined by a Welch’s t-test (* P < 0.05; ** P < 0.01; *** P < 0.001)

The same pattern was observed for the SON. Here again, the strongest decrease was observed in OXT neurons (Fig. 4D; *t* = 4.4; P = 0.002), and a smaller decrease was observed in AVP neurons (Fig. 4E; *t* = 2.9; P = 0.020). Similarly, the ratio between OXT and AVP neurons was decreased significantly in the SON as well (Fig. 4F; *t* = 3.0; P = 0.015).

## Discussion

In this study, we investigated the divergent differentiation of AVP and OXT MCNs using publicly available single-cell data of the developing mouse hypothalamus. Previous research on the differences between AVP and OXT MCNs has focused mainly on the neuron-specific expression of the respective neuropeptides, through investigation of specificity conferring promoter regions, rather than on the TF(s) responsible for the broader divergent differentiation. Instead, we leveraged single-cell computational analysis to investigate differentially active TFs between AVP and OXT MCN transcriptomes. Our analysis yielded five candidate TFs that may mediate the divergent differentiation of these neurons. For all candidates but RORA, we subsequently found diverging developmental gene expression patterns, supportive of the validity of the gene regulatory network analysis.

Our analysis of the SCRs of the *Avp* and *Oxt* promoters identified only RORA as a TF binding the *Avp* promoter, and three candidates (RORA, FOXP1, FOXP2) were predicted to bind the *Oxt* promoter. Interestingly, RORA was implicated here with several TFBSs in both promoters. However, these findings lack interpretability due to earlier conflicting findings and reports. The expression dynamics of *Rora* between populations were virtually identical, unlike the regulon activity and gene expression which were found enriched in *Avp* neurons. These findings are complicated by previous work, experimentally showing RORA to drive the mouse *Oxt* promoter^33^. In addition, RORA expression might not be evolutionarily conserved. In rats, previous work has found *Rora* to be enriched in *Oxt* neurons^34^, in contrast to our findings in mice. Due to this ambiguity surrounding the role of RORA, we focused our analysis on FOXP1 and FOXP2, instead.

Next, we used the *Foxp1*^+/-^ mouse model, the only model available to us. In this mouse line, we investigated whether *Foxp1* haploinsufficiency affects AVP and OXT neuron abundance in the PVN and SON. As this study focused solely on MCNs, it should be noted that the PVN contains both MCNs and parvocellular neurons (PCNs). Thus, the decreased OXT neuron abundance in the PVN may also be attributed to the PCNs present there, and this effect cannot be deconvolved with our current data. Nevertheless, the SON does contain only MCNs, and these results do conclusively determine the effect of *Foxp1* haploinsufficiency on MCNs.

We observed that the partial loss of *Foxp1* expression decreased abundance of both AVP and OXT neurons at P10. The effect on the AVP neurons was unexpected, though not without plausible explanation. Our modeled expression dynamics show decreased *Foxp1* and *Foxp2* expression in AVP neurons, but not entirely lacking. As such, these factors may affect AVP neuron development as well, albeit much less impactfully. Importantly, the OXT neuron abundance was disproportionally decreased at P10. This validates our prediction that there is a previously unrecognized role for FOXP1 in OXT neurons.

Although OXT and FOXP1 have not been related before in the literature, some notable phenotypic parallels can be found that connect the two. One of these is the mouse ultrasonic vocalizations (USVs) that are emitted by pups in various situations, including isolation from the nest. In mice that (partially) lack *Foxp1*, *Oxt* or its receptor (*Oxtr*), the number of USVs has been shown to decrease markedly^31,35–37^. Another parallel is autism spectrum disorder (ASD), a neurodevelopmental condition characterized by social deficits, repetitive behaviors, and sensory processing differences^38^. In mice, the knockouts of *Oxt* and *Oxtr* lead to social deficits^35,39^, while *Foxp1* knockout mice display both social deficits and repetitive behaviors^40^. In humans, ASD has been linked to lower plasma OXT levels and variants of the OXT receptor ^41,42^. Moreover, individuals with FOXP1 syndrome, harboring a deleterious variant in the *FOXP1* gene, often present with ASD as one of its various features^43^. Until now, these effects were often thought to occur due to the important role of FOXP1 in striatal medium spiny neurons^31^. However, given our new results, these phenotypic similarities may also partially be explained through the effect of FOXP1 variants on OXT MCNs.

This study does present with some limitations. First, based on our current data, we cannot conclusively determine if the deficient OXT MCN abundance in *Foxp1*^+/-^ mice should be viewed as delayed or impaired OXT MCN development, although functional implications may not differ strongly. Even delayed development could plausibly result in social deficits, due to the critical postnatal window for OXT system development^44^. This phenomenon can be observed in two mouse models with OXT dysfunction. In mice lacking the *Magel2* gene, a series of early-postnatal OXT injections rescued social and learning behavior in adulthood^45^. Similarly, in mice lacking the *Cntnap2* gene, postnatal OXT injections rescued the social behavioral phenotype in adults^46^. Interestingly, *Cntnap2* is a known target gene of FOXP1 and FOXP2^47,48^. These results show that the postnatal time period is crucial for normal development of OXT signaling, and *Foxp1*^+/-^ mice might be similarly affected even if OXT neuron development is only delayed.

Second, we used a whole-body mutant mouse line, and not a PVN- or MCN-specific knockout. Thus, the decreased neuron abundance observed in the mutant mice may have arisen through interactions with other brain regions or peripheral tissues. However, as our computational analysis independently suggests a role for FOXP1, we estimate this alternative explanation to be unlikely.

Third, as we did not test RORA, EBF3, BCL11B, or FOXP2 *in vivo*, we cannot rule out relevance of these other candidate TFs in differential differentiation. In particular, we might assume that FOXP2 likely contributes to OXT MCN development as well. This may be through FOXP1:FOXP2 heterodimerization, or FOXP2 may function as a redundancy mechanism. Considering the MCN phenotype of the haploinsufficient *Foxp1*^+/-^ mutant mice, the former seems the more likely option. For follow-up research, other viable candidates are the *Avp* MCN enriched TFs, which might drive AVPergic differentiation.

While there are many potential additional avenues for follow-up work to be tested, our data unequivocally shows that FOXP1 has a role in MCN development, and that its reduced presence disproportionally affects OXT neurons. This finding might help explain social aspects of FOXP1 syndrome and could unlock novel treatment strategies that target OXT system deficiencies.

## Methods

### scRNA-seq data collection and preprocessing

For the mouse developmental dataset from Romanov *et al.* (2020)^17^, sra data files were obtained from SRA BioProject PRJNA548917, using sra-toolkit 2.11.3. After prefetching the sra files, fastq files were derived with fasterq-dump command, using command-line flags “-p”, “-S”, “--include-technical”. Using the SRA metadata table, fastq files were merged per biological sample. For quantification of the count matrix, STAR^49^ 2.7.11b was used. First, a reference genome was created by running STAR with command-line flags “--runThreadN 20”, “--runMode genomeGenerate”. Files necessary to generate the reference genome were obtained from ENSEMBL. Then, the count matrix was computed for each sample, using STAR with command line flags “--runThreadN 16”, "--runDirPerm All_RWX", "--readFilesCommand zcat", "--outSAMtype None", "--soloType CB_UMI_Simple", "--soloCBwhitelist 737K-august-2016.txt", "-- soloBarcodeReadLength 0", "--soloCBlen 16", "--soloUMIstart 17", "--soloUMIlen 10", "-- soloStrand Forward", "--soloUMIdedup 1MM_CR", "--soloCBmatchWLtype 1MM_multi_Nbase_pseudocounts", "--soloUMIfiltering MultiGeneUMI_CR", "--soloCellFilter EmptyDrops_CR", "--clipAdapterType CellRanger4", "--outFilterScoreMin 30", "--soloFeatures Gene Velocyto", "--soloMultiMappers EM", "--outReadsUnmapped Fastx". Using python 3.11.4, scanpy 1.10.2 ^50^, R 4.4.2^51^, and Seurat 5.1.0^52^, the count matrices for the different samples were then loaded as single-cell dataset objects, and subsequently merged into a single dataset. Gene names were changes from ENSEMBL gene identifiers to MGI symbols. Finally, using original clustering metadata column “wtree” from Romanov *et al.* (2020)^17^, a subset was created to include only clusters 13, 14, 15, 16, 24, 26, 31, and 43. Of these, clusters 26 and 43 are the MCN clusters as defined in the original work. To enhance performance of the integration, the transcriptionally related PVN clusters 13-16, 24, and 31 were included in this subset.

### scRNA-seq data integration

To integrate the mouse developmental dataset, a subset of 1250 most highly variable genes was taken using the function highly_variable_genes, with arguments “n_top_genes = 1250” and “batch_key = sample”. Then, using the scvi-tools 1.2.0 model scVI^53^, the dataset was integrated on “sample”, using parameters “n_layers = 1”, “n_latent = 10”, “dropout_rate = 0.1”, “dispersion = gene”, “gene_likelihood = “nb”. The model was trained with arguments “max_epoch = 400”, “n_epochs_kl_warmup = 200”, “lr = 1e-2”, “lr_min = 1e-4”, “lr_patience = 33”, “lr_factor = 0.1**(1/3)”, “reduce_lr_on_plateau = True”, “lr_scheduler_metric = elbo_validation”, “check_val_every_n_epoch = 1”, “early_stopping = True”, early_stopping_patience= True”, and “early_stopping_monitor = elbo_validation”.

Subsequently, the trained scVI model was passed to scANVI^19^ and trained again, using the same parameters, except the additional or altered arguments “labels_key = Age”, “unlabeled_category = nan”, “max_epochs = 500”, “classification_ratio = 1.67”. The latent representation of the data was extracted used for downstream processing. Using original clustering metadata column “wtree” from Romanov *et al.* (2020)^17^, a subset was taken again, this time to only include MCN neurons (clusters 26 and 43).

The dataset was visualized with the RunUMAP function, using specified arguments “reduction = vae” and “dim = 1:10”. Neighborhoods were calculated with the FindNeighbors function, using the same arguments specified. Finally, the dataset was clustered using the FindClusters and FindSubClusters functions with resolution parameters set to 0.75 and 0.5, respectively. Finally, the resultant clusters were annotated.

### Gene regulatory network inference

Gene regulatory network activities were inferred with the pySCENIC pipeline. The pipeline was run with mostly default settings. For calculating the coexpression modules, the arboreto.algo.grnboost2 function was used with argument “seed = 0” specified, and the pyscenic.utils.modules_from_adjacencies function with argument “keep_only_activating = False” specified. Subsequently, for pruning the modules, pyscenic.prune.prune2df was used with arguments “rank_threshold = 1500”, “auc_threshold = 0.05” and “nes_threshold = 2.0” specified. The regulons were derived with the pyscenic.prune.df2regulons function, and regulon activity score were calculated with the pyscenic.aucell.aucell function, both with default arguments. For this section, python 3.10.13 was used with pyscenic 0.12.1^18^.

### Cell fate probability inference

To calculate cell fate probabilities, we used 3 modalities: RNA velocity, pseudotime and real-time using optimal transport analysis. For calculating RNA velocity, first a subset of highly variable genes was selected using function highly_variable_genes, with argument “flavor = seurat_v3”. Then, using VELOVI^23^, the RNA velocity matrix was calculated with default parameters. The training of the model used the same learning parameters as the scVI and scANVI training. The transition matrix was then extracted for downstream CellRank analysis, using default parameters. For calculating pseudotime, psupertime^24^ was used with regularization parameter “n_params = 30”, and ran with parameter “ordinal_data = Age”. Again, the transition matrix was extracted using default parameters. For the optimal-transport analysis, the moscot^25^ TemporalProblem class was used. The analysis was done with parameters “time_key = age_num”, “epsilon = 1e-3”, “tau_a = 0.95”, and “scale_cost = mean”. Here, the transition matrix was extracted using parameters “threshold = auto”, “self_transitions = all”, “conn_weight = 0.2”, and additional connection parameters “n_neighbors = 30”, “use_rep = X_pca”.

For the downstream CellRank 2^20^ analysis, the transition matrices were added with relative weight assigned: 30% pseudotime kernel, 15% velocity kernel, 55% real-time kernel. The GPCCA estimator was fit using parameters “cluster_key = Cluster”, n_states = [3,7]”, n_cells = 20”. Initial states were inferred with the predict_initial_states method. Terminal states were inferred with the predict_terminal_states method, using arguments “method = top_n”, and “n_states = 2”. Finally, fate probabilities were calculated with the compute_fate_probabilities method.

### Transcription factor binding site prediction

Sequences of the *Avp* and *Oxt* mouse promoters were retrieved from NCBI, using genome assembly GRCm39. From this, the SCR sequences were derived, resulting in the *Avp* SCR sequence “TCAACTATGATTTCCAGGTGACCCTCCAGTCGGCTCACCTCACTGATCGCACAGCACCAATCACTGTGGCAGTGGCTCCTGTCACACGGTGGCCGGTGACAGCCTGATGGCTGGCTCCCCTCCTCCACCACCCTCTGCACTGACAGGCCCACGTGTGTCCCCAGATGCCTGA” and *Oxt* SCR sequence “CCCCTTCCAGGCTGCTTCTCTTTTGAGTTCCAGGTCATTAGCAGAGACGATGACCTTGACCCTAGCCCAGACCCTGCAAATGAAGGGCCTGCCTCTAAACAGCGTGGAACAATTTG”. To predict TFBSs, the JASPAR^28^ position weight matrices (PWMs) used were: MA0071.1 (RORA), MA1637.2 (EBF3), MA0481.4 (FOXP1), MA0593.2 (FOXP2), and MA1989.2 (BCL11B). Using the TFBSTools 1.44.0 function searchSeq, SCR sequences were tested for TFBSs using the respective JASPAR PWMs. P-values computed were adjusted using the Holm-Bonferroni method.

### Animals

Mice were kept in a specific pathogen-free Biomedical Animal Facility under a 12-hour light/dark cycle with ad libitum access to water and food. All procedures were conducted in strict compliance with the National Institutes of Health Guidelines for the Care and Use of Laboratory Animals and approved by the National Institute of Mental Health animal care and use committee. The day of birth was considered as postnatal day (P) 0.5.

### Generation of *Foxp1*^+/-^ Animals

WT female mice were crossed with male mice, heterozygous for the *Foxp1* KO allele (*Foxp1*^+/-^)^29^. The *Foxp1*^+/-^ mice were backcrossed with C57BL/6J mice for at least 12 generations to obtain congenic animals.

### Brain perfusion and Immunohistochemistry

Mice were sacrificed at postnatal day 10 (P10), and brains were fixed overnight at 4°C in 4% paraformaldehyde (PFA). The following day, the brains were transferred to a 30% sucrose solution in PBS for cryoprotection. After approximately 48 hours, once the brains had sunk to the bottom of the well plates, they were gently blotted with absorbent paper, wrapped in aluminum foil, and stored at –20°C for at least 24 hours.

Cryosectioning was performed using a Leica cryostat. A total of 36 coronal sections, each 50 μm thick, were collected, starting from the anatomical level where the anterior commissures merged. All immunostaining procedures were carried out at room temperature using the free-floating method in well plates. Sections were first washed three times for 10 minutes each in PBS. They were then incubated for one hour in a blocking solution consisting of PBS with 2.5% normal donkey serum (ab7475), 2.5% normal goat serum (ab7481), and 0.1% Triton X-100. Primary antibody incubation was carried out overnight in the same blocking solution supplemented with mouse anti-Neurophysin 2 (1:1000, MABN856) and guinea pig anti-Oxytocin (1:500, 408 004, Synaptic Systems). Following incubation, sections were washed again three times for 10 minutes in PBS. Secondary antibody incubation was performed for four hours in PBS containing 0.1% Triton X-100, goat anti-guinea pig Alexa Fluor 488 (1:1000, A-11073), and donkey anti-mouse Alexa Fluor 594 (1:1000, ab150108). Sections were then washed three times for 10 minutes in PBS. The stained sections were mounted onto SuperFrost Plus adhesion slides (10149870, Epredia) and left to air-dry. VECTASHIELD Antifade Mounting Medium with DAPI (VEC-H-1200) was applied, and the slides were covered with glass coverslips (thickness no. 1, 101242, Marienfeld). Excess mounting medium was absorbed with paper, and nail hardener was applied around the edges of the coverslips to seal them.

### Slide Scanner Image Acquisition

All coronal brain sections (50 µm thick) containing the hypothalamic paraventricular (PVN), supraoptic (SON) and accessory nuclei (AN) were systematically collected from Foxp1-KO (n=7) and WT (n=7) mice. Brain sections (22-28 per animal) were immunofluorescence-stained for vasopressin and oxytocin as outlined in the previous section. Using an Olympus SLIDEVIEW VS200 system (Evident Scientific, Tokyo, Japan) equipped with a 20x objective, we acquired z-stacks (z-spacing 2.36 µm) of all ROIs containing AVP and OXT neurons in these sections. The system featured an S-Cite NOVEM 9-channell LED illumination system (Excelitas Technologies Corp., Massachusetts, USA), a fast pentaband filter wheel, and an iDS UI-3200SE-M-GL monochrome camera (IDS Imaging Development Systems GmbH, Obersulm, Germany). A standardized scanning protocol was employed across all samples to ensure consistent exposure and image resolution parameters (16 bit Grayscale, 0.35 µm/pixel, 6.45 ms exposure time at 455 nm for DAPI, 26.77 ms exposure time at 565 nm for Cy3. All scanned images were saved in VSI format for downstream analysis.

### Semi-automated quantification of oxytocin and vasopressin neurons

The image in .vsi format is opened using QuPath version 0.4.3. Brightness and contrast settings are adjusted using the "Min display" and "Max display" values to ensure that no image data is lost. Using the rectangle selection tool, the region of interest is selected, either the PVN or SON. In the case of the SON, two separate files are created to analyze the left and right sides independently. From the "Image" tab, the "Image commands" menu is opened and "Send region to ImageJ" is selected with the following settings: resolution set to 2, inclusion of ROI, inclusion of overlay, and all z-slices selected. Once the region is sent, ImageJ opens, and the image is saved via the "File" menu as a TIFF (.tif) file. The saved TIFF file is then opened in Imaris version 10.0.1. In the display adjustment settings, the red channel (channel 1) is turned of along with either the green (channel 2) or blue (channel 3) channel. Typically, the green channel is left on for initial analysis. The display intensity is manually adjusted to reduce background noise as much as possible, based on visual assessment rather than fixed values. The spot function is selected for cell detection. Spot creation parameters are left at their default settings. When needed, the "segment only a region of interest" option is used to simplify the selection of spots. All other algorithm settings remain unchanged. During the spot detection process, channel 2 (green) is first selected as the source, and the estimated cell diameter is set to 10.00 µm, while all other parameters for spot detection are left at their default values. The optional filtering step for spot selection is generally not used, but when cell density is high, the filter is used intuitively to limit false positives and reduce manual workload. The filter is applied conservatively to ensure only visually confirmed cells are selected. Following automatic detection, the "Edit" function is used to manually verify and adjust spot selection. This step is performed by visual inspection, rotating the 3D image to avoid missing any cells and to minimize human error, as no fixed parameters can be defined for this process. Once all cells are selected, the "Statistics" function is used to generate a count of the detected spots. The data is then exported using the "Export statistics on Tab display to file" function into an Excel (.xls) file. This process is repeated for each image containing a visible PVN, for both sides of the SON, and separately for each recording channel, including repeating the same analysis using the blue channel. All image processing, quantification, and data analysis were conducted entirely blind to experimental conditions.

### Image count data integration

Spot counts from individual Imaris reconstructions were aggregated per animal and cell type using R 4.4.2^51^.

### Confocal image acquisition

Representative high-resolution images for visualization purposes were acquired using a confocal microscope (Stellaris 5, Leica Microsystems, Wetzlar, Germany) equipped with a 40x objective (HC PL APO CS2 40×/1.25 GLYC, Leica Microsystems, Wetzlar, Germany). Image stacks were obtained from 50 μm-thick sections using a z-step interval of 1 μm. The acquisition settings were as follows: 1024 × 1024 pixels, 16-bit depth, pixel size 0.38 μm, and a zoom factor of 0.75

### Statistics

Statistical tests were performed in R 4.4.2^51^. P-values smaller than 0.05 were considered statistically significant. For the comparisons of neuron counts and OXT/AVP ratios between *Foxp1*^+/-^ and WT mice, a Welch’s t-test was performed.

### Data and code availability

The processed and annotated developmental mouse MCN dataset used for visualizations can be obtained from Zenodo^54^. The code used for scRNAseq analysis and visualization is publicly available at https://github.com/jberkh/2025_Avp_Oxt_Dif. The code used for the image count data processing is publicly available at https://github.com/tim-schubert/impro.

## References

1. Theofanopoulou C. Reconstructing the evolutionary history of the oxytocin and vasotocin receptor gene family: Insights on whole genome duplication scenarios. Dev Biol. 2021;479:99–106. doi:10.1016/j.ydbio.2021.07.012

2. Theofanopoulou C, Gedman G, Cahill JA, Boeckx C, Jarvis ED. Universal nomenclature for oxytocin–vasotocin ligand and receptor families. Nature. 2021;592(7856):747–755. doi:10.1038/s41586-020-03040-7

3. Boron WF, Boulpaep EL. Medical Physiology: A Cellular and Molecular Approach. Saunders Elsevier; 2012.

4. Am B, S T. The role of oxytocin in parturition. BJOG Int J Obstet Gynaecol. 2003;110 Suppl 20. doi:10.1016/s1470-0328(03)00024-7

5. Freund-mercier MJ, Moos F, Poulain DA, et al. Role of central oxytocin in the control of the milk ejection reflex. Brain Res Bull. 1988;20(6):737–741. doi:10.1016/0361-9230(88)90085-8

6. Kiyama H, Emson PC. Evidence for the Co-Expression of Oxytocin and Vasopressin Messenger Ribonucleic Acids in Magnocellular Neurosecretory Cells: Simultaneous Demonstration of Two Neurohypophysin Messenger Ribonucleic Acids by Hybridization Histochemistry. J Neuroendocrinol. 1990;2(3):257–259. doi:10.1111/j.1365-2826.1990.tb00401.x

7. Schonemann MD, Ryan AK, McEvilly RJ, et al. Development and survival of the endocrine hypothalamus and posterior pituitary gland requires the neuronal POU domain factor Brn-2. Genes Dev. 1995;9(24):3122–3135. doi:10.1101/gad.9.24.3122

8. Acampora D, Postiglione MP, Avantaggiato V, Di Bonito M, Simeone A. The role of Otx and Otp genes in brain development. Int J Dev Biol. 2000;44(6):669–677.

9. Wang W, Lufkin T. The Murine *Otp* Homeobox Gene Plays an Essential Role in the Specification of Neuronal Cell Lineages in the Developing Hypothalamus. Dev Biol. 2000;227(2):432–449. doi:10.1006/dbio.2000.9902

10. Hosoya T, Oda Y, Takahashi S, et al. Defective development of secretory neurones in the hypothalamus of Arnt2-knockout mice. Genes Cells. 2001;6(4):361–374. doi:10.1046/j.1365-2443.2001.00421.x

11. Michaud JL, DeRossi C, May NR, Holdener BC, Fan CM. ARNT2 acts as the dimerization partner of SIM1 for the development of the hypothalamus. Mech Dev. 2000;90(2):253–261. doi:10.1016/S0925-4773(99)00328-7

12. Michaud JL, Rosenquist T, May NR, Fan CM. Development of neuroendocrine lineages requires the bHLH–PAS transcription factor SIM1. Genes Dev. 1998;12(20):3264–3275. doi:10.1101/gad.12.20.3264

13. Xi D, Kusano K, Gainer H. Quantitative Analysis of Oxytocin and Vasopressin Messenger Ribonucleic Acids in Single Magnocellular Neurons Isolated from Supraoptic Nucleus of Rat Hypothalamus. Endocrinology. 1999;140(10):4677–4682. doi:10.1210/endo.140.10.7054

14. Glasgow E, Kusano K, Chin H, Mezey Ê, Young WS, Gainer H. Single Cell Reverse Transcription-Polymerase Chain Reaction Analysis of Rat Supraoptic Magnocellular Neurons: Neuropeptide Phenotypes and High Voltage-Gated Calcium Channel Subtypes. Endocrinology. 1999;140(11):5391–5401. doi:10.1210/endo.140.11.7136

15. Berkhout JB, Poormoghadam D, Yi C, Kalsbeek A, Meijer OC, Mahfouz A. An integrated single-cell RNA-seq atlas of the mouse hypothalamic paraventricular nucleus links transcriptomic and functional types. J Neuroendocrinol. 2024;36(2):e13367. doi:10.1111/jne.13367

16. Gainer H. Cell-type specific expression of oxytocin and vasopressin genes: An experimental odyssey. J Neuroendocrinol. 2012;24(4):528–538. doi:10.1111/j.1365-2826.2011.02236.x

17. Romanov RA, Tretiakov EO, Kastriti ME, et al. Molecular design of hypothalamus development. Nature. 2020;582(7811):246–252. doi:10.1038/s41586-020-2266-0

18. Van de Sande B, Flerin C, Davie K, et al. A scalable SCENIC workflow for single-cell gene regulatory network analysis. Nat Protoc. 2020;15(7):2247–2276. doi:10.1038/s41596-020-0336-2

19. Xu C, Lopez R, Mehlman E, Regier J, Jordan MI, Yosef N. Probabilistic harmonization and annotation of single-cell transcriptomics data with deep generative models. Mol Syst Biol. 2021;17(1):e9620. doi:10.15252/msb.20209620

20. Weiler P, Lange M, Klein M, Pe’er D, Theis F. CellRank 2: unified fate mapping in multiview single-cell data. Nat Methods. Published online June 13, 2024. doi:10.1038/s41592-024-02303-9

21. Li S, Weidenfeld J, Morrisey EE. Transcriptional and DNA binding activity of the Foxp1/2/4 family is modulated by heterotypic and homotypic protein interactions. Mol Cell Biol. 2004;24(2):809–822. doi:10.1128/MCB.24.2.809-822.2004

22. Sin C, Li H, Crawford DA. Transcriptional Regulation by FOXP1, FOXP2, and FOXP4 Dimerization. J Mol Neurosci. 2015;55(2):437–448. doi:10.1007/s12031-014-0359-7

23. Gayoso A, Weiler P, Lotfollahi M, et al. Deep generative modeling of transcriptional dynamics for RNA velocity analysis in single cells. Nat Methods. 2024;21(1):50–59. doi:10.1038/s41592-023-01994-w

24. Macnair W, Gupta R, Claassen M. psupertime: supervised pseudotime analysis for time- series single-cell RNA-seq data. Bioinformatics. 2022;38(Supplement_1):i290–i298. doi:10.1093/bioinformatics/btac227

25. Schiebinger G, Shu J, Tabaka M, et al. Optimal-Transport Analysis of Single-Cell Gene Expression Identifies Developmental Trajectories in Reprogramming. Cell. 2019;176(4):928–943.e22. 10.1016/j.cell.2019.01.006

26. Ponzio TA, Fields RL, Rashid OM, Salinas YD, Lubelski D, Gainer H. Cell-type specific expression of the vasopressin gene analyzed by AAV mediated gene delivery of promoter deletion constructs into the rat SON in vivo. PloS One. 2012;7(11):e48860. doi:10.1371/journal.pone.0048860

27. Fields RL, Ponzio TA, Kawasaki M, Gainer H. Cell-Type Specific Oxytocin Gene Expression from AAV Delivered Promoter Deletion Constructs into the Rat Supraoptic Nucleus in vivo. PLOS ONE. 2012;7(2):e32085. doi:10.1371/journal.pone.0032085

28. Rauluseviciute I, Riudavets-Puig R, Blanc-Mathieu R, et al. JASPAR 2024: 20th anniversary of the open-access database of transcription factor binding profiles. Nucleic Acids Res. 2024;52(D1):D174–D182. doi:10.1093/nar/gkad1059

29. Wang B, Weidenfeld J, Lu MM, et al. Foxp1 regulates cardiac outflow tract, endocardial cushion morphogenesis and myocyte proliferation and maturation. Dev Camb Engl. 2004;131(18):4477–4487. doi:10.1242/dev.01287

30. Wang J, Rappold GA, Fröhlich H. Disrupted Mitochondrial Network Drives Deficits of Learning and Memory in a Mouse Model of FOXP1 Haploinsufficiency. Genes. 2022;13(1):127. doi:10.3390/genes13010127

31. Araujo DJ, Anderson AG, Berto S, et al. FoxP1 orchestration of ASD-relevant signaling pathways in the striatum. Genes Dev. 2015;29(20):2081–2096. doi:10.1101/gad.267989.115

32. Althammer F, Wimmer MC, Krabichler Q, et al. Analysis of the hypothalamic oxytocin system and oxytocin receptor-expressing astrocytes in a mouse model of Prader–Willi syndrome. J Neuroendocrinol. 2022;34(12):e13217. doi:10.1111/jne.13217

33. Chu K, Zingg HH. Activation of the mouse oxytocin promoter by the orphan receptor RORalpha. J Mol Endocrinol. 1999;23(3):337–346. doi:10.1677/jme.0.0230337

34. Humerick M, Hanson J, Rodriguez-Canales J, et al. Analysis of Transcription Factor mRNAs in Identified Oxytocin and Vasopressin Magnocellular Neurons Isolated by Laser Capture Microdissection. PLOS ONE. 2013;8(7):e69407. doi:10.1371/journal.pone.0069407

35. Takayanagi Y, Yoshida M, Bielsky IF, et al. Pervasive social deficits, but normal parturition, in oxytocin receptor-deficient mice. Proc Natl Acad Sci. 2005;102(44):16096–16101. doi:10.1073/pnas.0505312102

36. Fröhlich H, Rafiullah R, Schmitt N, Abele S, Rappold GA. Foxp1 expression is essential for sex-specific murine neonatal ultrasonic vocalization. Hum Mol Genet. 2017;26(8):1511–1521. doi:10.1093/hmg/ddx055

37. Winslow JT, Hearn EF, Ferguson J, Young LJ, Matzuk MM, Insel TR. Infant Vocalization, Adult Aggression, and Fear Behavior of an Oxytocin Null Mutant Mouse. Horm Behav. 2000;37(2):145–155. doi:10.1006/hbeh.1999.1566

38. Lord C, Brugha TS, Charman T, et al. Autism spectrum disorder. Nat Rev Dis Primer. 2020;6(1):1–23. doi:10.1038/s41572-019-0138-4

39. Winslow JT, Insel TR. The social deficits of the oxytocin knockout mouse. Neuropeptides. 2002;36(2):221–229. doi:10.1054/npep.2002.0909

40. Bacon C, Schneider M, Le Magueresse C, et al. Brain-specific Foxp1 deletion impairs neuronal development and causes autistic-like behaviour. Mol Psychiatry. 2015;20(5):632–639. doi:10.1038/mp.2014.116

41. LoParo D, Waldman ID. The oxytocin receptor gene (OXTR) is associated with autism spectrum disorder: a meta-analysis. Mol Psychiatry. 2015;20(5):640–646. doi:10.1038/mp.2014.77

42. John S, Jaeggi AV. Oxytocin levels tend to be lower in autistic children: A meta-analysis of 31 studies. Autism Int J Res Pract. 2021;25(8):2152–2161. doi:10.1177/13623613211034375

43. Bacon C, Rappold GA. The distinct and overlapping phenotypic spectra of FOXP1 and FOXP2 in cognitive disorders. Hum Genet. 2012;131(11):1687–1698. doi:10.1007/s00439-012-1193-z

44. Muscatelli F, Matarazzo V, Chini B. Neonatal oxytocin gives the tempo of social and feeding behaviors. Front Mol Neurosci. 2022;15. doi:10.3389/fnmol.2022.1071719

45. Meziane H, Schaller F, Bauer S, et al. An Early Postnatal Oxytocin Treatment Prevents Social and Learning Deficits in Adult Mice Deficient for *Magel2*, a Gene Involved in Prader-Willi Syndrome and Autism. Biol Psychiatry. 2015;78(2):85–94. doi:10.1016/j.biopsych.2014.11.010

46. Peñagarikano O, Lázaro MT, Lu XH, et al. Exogenous and evoked oxytocin restores social behavior in the Cntnap2 mouse model of autism. Sci Transl Med. 2015;7(271):271ra8- 271ra8. doi:10.1126/scitranslmed.3010257

47. O’Roak BJ, Deriziotis P, Lee C, et al. Exome sequencing in sporadic autism spectrum disorders identifies severe de novo mutations. Nat Genet. 2011;43(6):585–589. doi:10.1038/ng.835

48. Vernes SC, Newbury DF, Abrahams BS, et al. A Functional Genetic Link between Distinct Developmental Language Disorders. N Engl J Med. 2008;359(22):2337–2345. doi:10.1056/NEJMoa0802828

49. Dobin A, Davis CA, Schlesinger F, et al. STAR: ultrafast universal RNA-seq aligner. Bioinformatics. 2013;29(1):15–21. doi:10.1093/bioinformatics/bts635

50. Wolf FA, Angerer P, Theis FJ. SCANPY: large-scale single-cell gene expression data analysis. Genome Biol. 2018;19(1):15. doi:10.1186/s13059-017-1382-0

51. R Core Team. R: A Language and Environment for Statistical Computing. Published online 2024.

52. Hao Y, Stuart T, Kowalski MH, et al. Dictionary learning for integrative, multimodal and scalable single-cell analysis. Nat Biotechnol. 2024;42(2):293–304. doi:10.1038/s41587-023-01767-y

53. Lopez R, Regier J, Cole MB, Jordan MI, Yosef N. Deep generative modeling for single-cell transcriptomics. Nat Methods. 2018;15(12):1053–1058. doi:10.1038/s41592-018-0229-2

54. Berkhout JB. FOXP1 differentially regulates the development of murine vasopressin and oxytocin magnocellular neurons. Published online June 4, 2025. doi:10.5281/zenodo.15594918

